# Internalization of myelin debris by neutrophils fuels inflammation

**DOI:** 10.64898/2026.04.30.722078

**Authors:** Mirre De Bondt, Janne Renders, Yana Sterckx, Sofie Vandendriessche, Jana Van Broeckhoven, Annet van de Waterweg Berends, Noëmie Pörtner, Lotte Vanbrabant, Vívian Louise Soares de Oliveira, Jeroen Bogie, Sam Vanherle, Pedro Elias Marques, Niels Hellings, Sofie Struyf

## Abstract

Progressive neurodegeneration in the central nervous system (CNS) in multiple sclerosis (MS) is driven by chronic inflammatory demyelination. Neutrophils are increasingly recognized as versatile innate immune cells with potentially underappreciated roles in CNS inflammation, but their contribution to MS pathology remains poorly understood. Interestingly, we observed foamy neutrophils in active CNS lesions of MS patients. Therefore, we investigated the ability of human neutrophils to internalize myelin debris and assessed how this impacts their functional phenotype. Neutrophils exhibited efficient myelin uptake, peaking between 3 and 6 hours, predominantly through complement opsonization and internalization via complement receptor 3. Prolonged exposure to high concentrations of myelin induced a pro-inflammatory phenotype, marked by increased production of reactive oxygen species, neutrophil extracellular traps, and inflammatory mediators such as CXCL8 and CCL3. Gene expression analysis revealed a dose-dependent inflammatory signature after myelin uptake, characterized by gradual upregulation of *CXCL8* and decreased *ARG1* expression, suggesting a shift toward a pro-inflammatory neutrophil phenotype. These findings provide novel insights into the role of neutrophils in myelin clearance and inflammation in the CNS, highlighting complement receptor 3-mediated uptake and downstream pro-inflammatory activation as key mechanisms.

## 1. Introduction

Demyelinating diseases comprise a group of disorders where the myelin sheath around neurons is damaged, hampering transduction of action potentials along the axons. Demyelination is apparent in many diseases affecting the central nervous system (CNS), with multiple sclerosis (MS) being the most common (1). The chronic nature of inflammation in such diseases leads to irreversible nerve damage and disability with no cure available yet. Early in disease pathology, spontaneous remyelination allows repair of the damaged axons (2). Remyelination is initiated by the recruitment and differentiation of oligodendrocyte precursor cells (OPCs), that will synthetize new myelin components (3). Furthermore, innate immune cells such as macrophages and microglia contribute to this process by releasing OPC-differentiating factors and clearing myelin debris, the accumulation of which has been shown to inhibit remyelination (4–6). Additionally, it has been shown that temporary myelin uptake induces a disease-resolving phenotype, while prolonged intracellular accumulation of myelin skews these phagocytes to a pro-inflammatory phenotype (7,8). In later disease stages, chronic demyelination combined with age-dependent declines in intrinsic remyelination capacity hinders myelin regeneration and disease remission. Consequently, remyelinating areas are left vulnerable resulting in a slowly expanding MS lesions, now commonly named “smouldering” MS lesions, where a rim of active myeloid cells surrounds a hypocellular core and attacks the surrounding myelin (9).

Alongside microglia and macrophages, neutrophils are reported to take up myelin debris in both the cuprizone and experimental autoimmune encephalomyelitis (EAE) models of MS (10,11). Despite extensive evidence for involvement of these innate immune cells from both animal models and patient samples (12–14), neutrophils are understudied in MS research (15). Being professional phagocytes, neutrophils contain a set of specialized receptors that mediate recognition and uptake of targets, such as pattern recognition receptors, integrins (including complement receptors) and Fc receptors. They can internalize both opsonized and non-opsonized targets, however, opsonization maximizes phagocytic efficiency. Neutrophils express two main opsonin receptors: the Fc receptors and the β2-integrin complement receptors.

Myelin debris is susceptible to opsonization by either autoantibodies or activated complement components (e.g., C3b) (16). Evidence for complement involvement in demyelinating disorders has been provided in the EAE mouse model for MS; proteomic analyses showed upregulation of C3 and C1q, and C3 knockout mice were protected from synapse loss (17,18). In the cuprizone mouse model for demyelination, complement deposition was detected in the corpus callosum and overexpression of C3a and C5a resulted in augmented demyelination, accompanied by microglia activation and recruitment of neutrophils (19,20). Here, we sought to define the presence of myelin-containing foamy neutrophils in postmortem human MS lesions, elucidate the molecular mechanisms driving myelin internalization, and determine the impact of intracellular myelin accumulation on their inflammatory phenotype.

Our findings indicate that neutrophils are present within demyelinating CNS lesions of EAE and MS tissue, and accumulate myelin-loaded lipid droplets *in vitro.* Myelin uptake is mediated through opsonization with soluble complement factors and binding to complement receptor 3. Furthermore, chronic exposure of neutrophils to high-dose myelin drives a pro-inflammatory shift in their phenotype and functional state, characterized by an increased release of reactive oxygen species, neutrophil extracellular traps and inflammatory enzymes, proteases and mediators.

## 2. Materials and methods

### 2.1 Immunofluorescent staining of MS lesions

Snap-frozen brain tissue containing active MS lesions was obtained from the Netherlands Brain Bank (NBB, Amsterdam, Netherlands). Cryostat sections (10 µm) were prepared from these lesions and first incubated for 10 min in cold acetone (-20°C) and afterwards for 5 min in 70% ethanol at -20°C. Sections were washed twice for 3 min with PBS-T (PBS + 0.05% Tween-20) at RT and blocked with 10% Dako protein block (Agilent, Santa Clara, CA, USA) in PBS-T for 30 min at RT. Afterwards, samples were incubated overnight at 4°C with primary antibodies; mouse anti hu CD66b (BD Biosciences, Franklin Lakes, NJ, USA; #555723, 1:100 in PBS) and rabbit anti human PLIN-2 (Abcam, Cambridge, UIK; #108323, 1:100 in PBS). Following incubation, samples were washed 7 times for 2 min with PBS-T and incubated for 40 min at RT with secondary antibodies (DαRb555, #A31570 and DαMs488, #A11055; both from Invitrogen, Carlsbad, CA, USA) 1:400 in PBS. Sections were washed 3 times 5 min with PBS and incubated for 10 min at RT with DAPI (1:10.000 in PBS; Invitrogen). Afterwards, samples were washed again 3 times for 5 min with PBS and incubated for 15 min at RT in 0.1% Sudan Black (diluted in 70% ethanol) and rinsed with PBS after which they were mounted on slides. Lesions were imaged using the LSM900 confocal microscope (Zeiss).

### 2.2 DAB staining of MS lesions

The brain sections were first incubated for 10 min in cold acetone (-20°C) and afterwards dried for 30 min at RT. Sections were washed 3 times for 3 min with PBS at RT and blocked with Dako protein block for 30 min at RT. Afterwards, samples were incubated overnight at 4°C with primary antibodies (HLA DR; Thermo Fisher Scientific, Waltham, MA, USA, #MA1-90486 and PLP; Bio-Rad, Hercules, CA, USA, #MCA839G) both diluted 1:100 in 10% Dako protein block in PBS. Following incubation, samples were washed 3 times 5 min with PBS and incubated for 30 min at RT with Dako EnVision-HRP (Agilent, #K4065). Sections were washed 3 times 5 min with PBS, stained with 3,3′-diaminobenzidine (DAB) for 1 minute and washed 3 times with water. Samples were then stained with haematoxylin for 5 min at RT and rinsed with water. Next, slides were dehydrated by incubating them for 3 minutes in each of the following ethanol solutions: 70% - 80% - 95% - 100% - 100%. Afterwards, slides were incubated for 5 minutes in Xylene I and Xylene II after which cover slips were added using DPX mounting. Lesions were imaged using the Axioscan (Zeiss, Oberkochen, Germany).

### 2.3 Sample collection and neutrophil purification

Blood and serum samples from healthy volunteers were collected using EDTA-coated and thrombin-coated vacutainer blood collection tubes, respectively (BD Biosciences). For serum sampling, collected blood was incubated for 30 min at room temperature (RT) and centrifuged for 10 min at 1500 g. A part of the serum samples was heated at 56°C for 30 min (heat-inactivation, HI) to inactivate the complement components. Neutrophils were purified within 15 min after blood withdrawal using the EasySep human neutrophil isolation kit (STEMCELL technologies, Vancouver, Canada), according to the manufacturer’s protocol. This study was approved by the Ethics Committee of the University Hospital Leuven (study number: S58418).

### 2.4 Myelin debris labelling and opsonization

Purified myelin from *postmortem* healthy human brain tissue was isolated as previously described (21). Myelin was dissolved in 0.1 M sodium bicarbonate buffer (pH 8.5) and disintegrated into debris with a pellet mixer (VWR Avantor, Haasrode, Belgium) for 5 min. After centrifugation for 5 min (17000 g), the supernatant was discarded, and the pellet was resuspended in sodium bicarbonate buffer. Myelin debris was labelled with pHrodo succinimidyl ester (0.02 μL/μg myelin; Invitrogen) for 30 min at RT. Excess pHrodo was washed away twice with phosphate-buffered saline (PBS). Labelled myelin debris was then opsonized with 20% human serum, 20% HI serum, 20% C3- or C1q-depleted serum (Complement technologies, Tyler, TX, USA) or PBS for 1 h at 37°C. Afterwards, myelin was centrifuged (17000 g) for 5 min and washed twice for removal of unbound fragments. The pellet was finally dissolved in appropriate medium for subsequent assays.

### 2.5 Internalization of myelin by neutrophils

#### 2.5.1 IncuCyte analysis

Neutrophils were plated in a 48-well plate (5 x 10^4^ cells/well) and incubated for 30 min with Calcein AM (1 μM; Invitrogen) to stain for cell viability (22). For receptor-blocking experiments, latrunculin B (10 μg/mL; Sigma-Aldrich, St. Louis, MO, USA) or one of the following antibodies were added to the neutrophils: anti-CD11b (BioLegend, San Diego, CA, USA; #101201, 10 µg/mL), isotype control (BioLegend, #400601, 10 µg/mL) or human polyclonal IgG (100 µg/mL, freshly purified). After incubation (15 min at 37°C), PBS or a mix of cytokines (50 ng/mL TNF-α, 10 ng/mL IFN-γ and 10 ng/mL IL-1β; all from PeproTech, Cranbury, NJ, USA), and opsonized pHrodo-labelled myelin debris (1, 10 or 100 µg/mL) were added to the cells. Plates were imaged every hour in the IncuCyte S3 system (Sartorius, Göttingen, Germany) for 24 h and analyzed with IncuCyte analysis software (Sartorius). In every well, the number of phagocytosing events (i.e., the number of calcein-green and pHrodo-red fluorescent overlapping events) was determined for all acquired images. This number was normalized to the initial count of viable neutrophils in the corresponding well. The percentage of phagocytosis was plotted over time, and the maximal percentage of phagocytosis was used to compare different experimental conditions.

#### 2.5.2 Confocal microscopy

Myelin debris internalization by neutrophils was verified using confocal microscopy. Therefore, neutrophils were pre-treated with Calcein AM (1 μM) for 30 min at 37°C and plated (1.5 x 10^4^ cells/well) in a Lab-Tek chamber (Ibidi, Gräfelfing, Germany) in HBSS buffer (containing Ca^2+^ and Mg^2+^; VWR Avantor). Myelin debris was opsonized and labelled, as described above, and dissolved in HBSS buffer. Neutrophils were then incubated with myelin debris for 3 h at 37°C. Afterwards, cells were imaged using the dragonfly spinning-disk confocal microscope (Andor Technology, Belfast, Ireland) with 63x objective and analyzed using Imaris Software 9.8.0 (Andor Technology). For lipid droplet staining, neutrophils were first fixed after 3 h of myelin phagocytosis with 4% paraformaldehyde in HBSS buffer for 30 min at RT, after which they were washed with HBSS buffer. Next, the cells were permeabilized with 0.1% Triton X-100 for 15 min at RT and washed again with HBSS. The cells were then stained with Hoechst (10 µg/mL) and Phalloidin-647 (1:400) in combination with 300 nM Nile Red or 2 µm Bodipy (all from Thermo Fisher Scientific) for 1 h at RT, washed afterwards and imaged.

### 2.6 SDS-PAGE and Western blot

For low molecular weight gel electrophoresis, samples were diluted in reducing loading buffer (50% glycerol, 11.5% SDS, 0.1% bromophenol blue, 25% β-mercaptoethanol, pH 6.8), incubated for 10 min at 95°C and loaded onto a pre-cast 12% tris-glycine mini gel (Invitrogen) for protein separation. Samples for high molecular weight gel electrophoresis were diluted in non-reducing loading buffer (50% glycerol, 11.5% SDS, 0.1% bromophenol blue) and loaded onto a pre-cast 6% tris-glycine mini gel (Invitrogen). Protein samples and low molecular weight protein ladder (Precision Plus Protein Standard; Bio-Rad) or high molecular weight protein ladder (Spectra High Range Multicolor Protein Ladder; Thermo Fisher Scientific) were loaded into the gel. Separation was obtained through application of a constant current and 200 V for 30 min in running buffer (192 mM glycine, 25 mM Tris and 0.1% SDS). After separation, the gels were transferred to a PVDF membrane (Bio-Rad) in a Turbo blot machine (Bio-Rad) for 30 min at 25 V. Afterwards, the membrane was blocked for 1 h at RT in block buffer, containing 5% non-fat dry milk (Bio-Rad) in TBS wash buffer (20 mM Tris, 150 mM NaCl, 0.01% tween20, pH 7.5). The membrane was washed for 8 min in wash buffer, and the primary anti-C3/C3b antibody (Bio-Rad; #MCA6151GA, 1/500 diluted in block buffer) was added to the membrane, followed by overnight incubation at 4°C. The membrane was washed 3 times for 8 min with wash buffer and incubated with a goat anti-mouse horseradish peroxidase-labelled antibody (Jackson Immunoresearch, Cambridgeshire, UK; 1/10.000 diluted in block buffer) for 1 h at RT. Afterwards, the membrane was washed 3 times for 8 min with wash buffer and visualized with chemiluminescence substrate (Super Signal West Pico PLUS, Thermo Fisher Scientific) and analyzed with the Fusion Solo S (Vilber, Marne-la-Vallée, France).

### 2.7 Reactive oxygen species (ROS) quantification

Neutrophils were incubated with buffer or different doses of unlabeled opsonized myelin debris (1, 10 or 100 µg/mL) for 6 or 24 h. Afterwards, cells were harvested, pelleted and resuspended in RPMI medium for counting. Neutrophils (1.5 x 10^5^ cells/well) were added to a white luminescence plate (Perkin Elmer, Waltham, MA, USA) and stimulated with PBS, PMA (Sigma-Aldrich, 150 ng/mL) or a combination of TNF-α priming for 10 min (50 ng/mL) and subsequent IL-1β stimulation (50 ng/mL). Luminol (Sigma-Aldrich, 2 mM diluted in RPMI) was added to all conditions and chemiluminescence was measured for 3 h using a Clariostar plate reader (BMG Labtech, Ortenberg, Germany).

### 2.8 NETosis assay

Purified neutrophils were labelled with SYTOX Green (50 nM in RPMI, Invitrogen), transferred to a black, clear-bottom 96-well plate (5 x 10^4^ cells/well) and incubated for 15 min at 37°C to allow cell adherence. Afterwards, serum-opsonized myelin debris (1, 10 or 100 µg/mL) was carefully added, and cells were imaged every hour for 10 h at 37°C in the IncuCyte imaging system. The relative area of SYTOX green fluorescence (released DNA) was determined using the IncuCyte S3 software and normalized to the cellular area at timepoint 0 h.

### 2.9 Interleukin-8 (IL-8/CXCL8), chemokine (C-C motif) ligand 3 (CCL3/MIP-1α) and neutrophil elastase (NE) ELISA

Neutrophils were incubated with buffer or different concentrations of serum-opsonized myelin debris (1, 10 or 100 µg/mL) for 2 h in RPMI-1640 (for NE) or 24 h in induction medium (RPMI-1640 with phenol red + 10% FCS + 1/1000 gentamycin + 5 ng/ml GM-CSF; for CXCL8 and CCL3). Afterwards, supernatants were harvested, centrifuged for 5 min at 17000 g to remove cells and debris, and stored at -20°C for cytokine measurement by ELISA. The concentration of CXCL8 was measured by an in-house developed sandwich ELISA (23). The concentration of CCL3 or NE was determined with the human CCL3/MIP-1α or NE DuoSet (R&D systems, Minneapolis, MN, USA; P139302 and #DY9167-05), according to the manufacturer’s protocol.

### 2.10 Gelatin zymography for matrix metalloproteinase (MMP) quantification

To assess protein levels of the gelatinases (pro-)MMP-2, and (pro-)MMP-9, in gel gelatin zymography was performed as previously described (24). Gels were cast with a 7.5% acrylamide resolving gel containing 1 mg/mL gelatin (Sigma-Aldrich), overlaid with a 5% stacking gel. Following polymerization, gels were loaded into an electrophoresis system containing running buffer (25 mM Tris, 192 mM glycine, 0.1% SDS). Samples were prepared in non-reducing loading buffer (125 mM Tris-HCl, pH 6.8, 4% SDS, 20% glycerol, 0.1% bromophenol blue) and loaded onto the gel. After electrophoretic protein separation, gels were washed twice for 20 min in reactivation buffer (2.5% Triton X-100). Gels were then incubated overnight in 10 mM CaCl_2_ and 50 mM Tris–HCl, pH 7.5 at 37°C, to allow enzymatic activity. Proteolytic activity was visualized by staining the gels with 0.1% Coomassie Brilliant Blue R-350 (GE Healthcare), and zymograms were quantified using ImageQuant TL software (GE Healthcare). Molecular weights were determined using a reference standard consisting of trimeric pro-MMP-9, monomeric pro-MMP-9, and a low-molecular weight pro-MMP-9 domain deletion mutant lacking the O-glycosylated and hemopexin domains (proMMP-9 ΔOGΔHem).

### 2.11 Gene expression quantification

#### 2.11.1 mRNA isolation

Neutrophils were incubated with buffer or different concentrations of serum-opsonized myelin debris (1, 10 or 100 µg/mL) for 6 or 12 h in RPMI buffer. Afterwards, neutrophils (4 x 10^6^ per condition) were harvested, centrifuged for 5 min at 300 g, and washed once with PBS. Cells were then lysed with lysis buffer (1% β-mercaptoethanol in RLT buffer) and homogenized with sterile syringes and needles. Subsequently, mRNA was isolated using the RNeasy mini kit with on column DNase treatment (Qiagen, Hilden, Germany), according to the manufacturer’s protocol. The mRNA concentration and quality were determined at 260 nm using a NanoDrop ND-1000 spectrophotometer (Isogen Life Science, Utrecht, The Netherlands).

#### 2.11.2 cDNA synthesis

Next, mRNA was converted to cDNA by reverse transcription. The following master mix was prepared: RT buffer, nucleotides, random primers and RT enzyme (all diluted to 1X in RNase-free water; Applied Biosystems, Foster City, CA, USA). Equal volumes of mRNA sample (500 ng) and master mix were combined to perform an RT reaction. As a direct control for the reaction, an RT negative sample was prepared, containing all components of the master mix except for RT enzyme. The cDNA synthesis was performed in a thermocycler (Bio-Rad) using the following program: 10 min (25°C), 120 min (37°C), 5 min (85°C) and kept stable at 4°C. Afterwards, synthesized cDNA was stored at -20°C.

#### 2.11.3 Quantitative polymerase chain reaction (qPCR/RT-PCR)

A MicroAmp 96-well plate (Thermo Fisher Scientific) was loaded with 20 ng cDNA sample and 1X primer/probe mix (see table 1; all from Integrated DNA Technologies, Leuven, Belgium), diluted in Taqman mix (Applied Biosystems). The QuantStudio real-time PCR system (Applied Biosystems) was used for the following qPCR reaction: 2 min (50°C), 10 min (95°C) followed by 40 cycles of 15 s (95°C) and 1 min (60°C). The housekeeping gene 18S was used as positive control, and an RT negative sample and RNase free water were used as negative controls. For analysis, the standard ΔΔCt method was used to calculate fold changes from cycle threshold values (25).

**Table 1.** Primers/probe pairs used for qPCRa.

| Target | Lot | Fluorophore |
| --- | --- | --- |
| 18S | Hs.PT.39a.22214856.g | HEX |
| *Arginase-1(ARG1)* | Hs.PT.56a.20779559 | FAM |
| *Heme oxygenase 1 (HMOX1)* | Hs.PT.58.45340055 | HEX |
| *Interleukin 8 (CXCL8)* | IL-8 set 3 assay | FAM |
| *Transforming growth factor-β (TGFB1)* | Hs.PT.58.39813975 | Cy5 |
| *Tumor necrosis factor-α (TNF)* | Hs.PT.51.15510722 | HEX |
| *Vascular endothelial growth factor (VEGF)* | Hs.PT.49a.20806656 | FAM |

<sup>a</sup> All ordered from Integrated DNA Technologies

#### 2.11.4 Bulk RNA sequencing

Based on pilot experiments testing 0h, 3h, 6h and 12h incubation periods we selected 12h incubation time as this showed the strongest upregulation of CXCL8 gene expression. Neutrophils were incubated with PBS or myelin debris (1 or 100 µg/mL) and mRNA was isolated as described above. RNA sequencing was performed by Novogene (Munich, Germany). Bioinformatic processing and quality control, also conducted by Novogene, yielded 90–94% total mapping rates and 95–98% clean reads. Differential gene expression analysis was performed using DESeq2, applying a threshold of |log2(FoldChange)|≥ 1 and an adjusted p-value ≤ 0.05. Hierarchical clustering was conducted using log2(FPKM+1) values. Downstream functional enrichment and pathway analyses were carried out using g:Profiler and STRING (search tool for the retrieval of interacting genes/proteins).

### 2.12 Statistics

The normality of all data was first evaluated using a Shapiro-Wilk test and subsequently corresponding downstream statistical testing was applied with a correction for multiple testing. The applied statistical tests are indicated in the figure legends. Statistical analysis and visualization of the data were performed using GraphPad Prism 9.3.1.

## Results

### 3.1 Neutrophils in MS and EAE lesions actively internalize myelin

To investigate whether neutrophils are present in demyelinating CNS lesions, we analyzed spinal cord tissue from mice with experimental autoimmune encephalomyelitis (EAE) and postmortem brain tissue from patients with multiple sclerosis (MS). Neutrophil infiltration was detected in regions of demyelination in both EAE and MS (**Supplementary fig 1; Fig 1A**). To further characterize their phenotype in human disease, we stained MS tissue for CD66b (neutrophils) and perilipin-2 (PLIN2), a lipid droplet-associated protein enriched in foamy phagocytes. This revealed the presence of lipid-loaded neutrophils within actively demyelinating, immune-cell rich lesion areas (**Fig 1A**).

**Figure 1.**
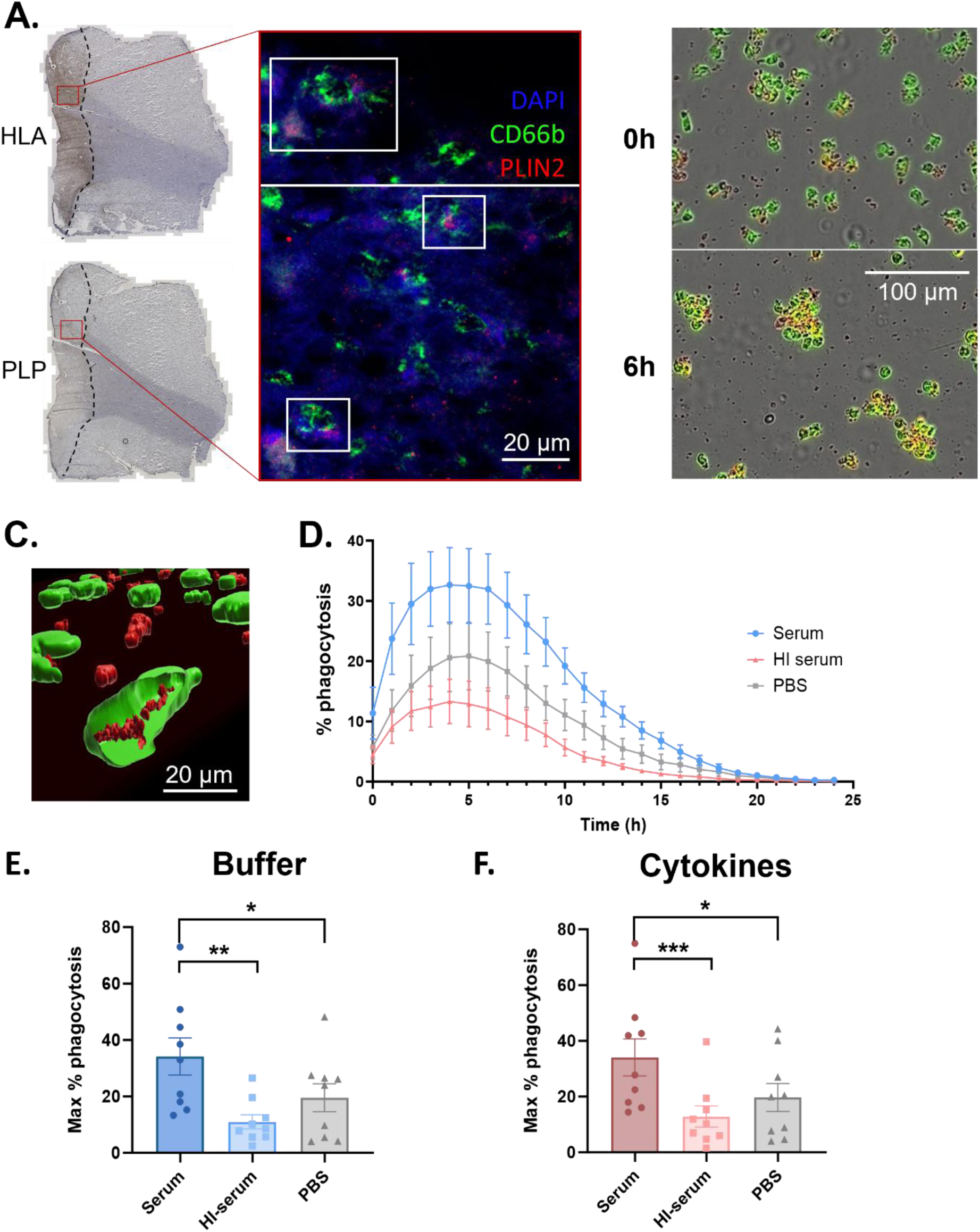
Human neutrophils take up myelin debris. **(A)** Human postmortem MS brain tissue was stained for HLA and PLP to delineate regions of active immune cell infiltration and demyelination. Lesions were stained for immunofluorescence with DAPI, CD66b (neutrophils) and PLIN2 (lipid droplets) to show myelin accumulation within neutrophils. **(B-F)** Calcein-labelled purified neutrophils from healthy donors were incubated with pHrodo-labelled myelin debris (75 µg/mL), whereby fluorescent images were generated every hour for 24 h with the IncuCyte imaging system. **(B)** Representative IncuCyte images of serum-opsonized myelin debris (red) phagocytosis by human neutrophils (green) at time points 0 h and 6 h. **(C)** Cross-section of human calcein-labelled neutrophils (green) with ingestion of pHrodo-labelled myelin debris (red) by confocal imaging (63x objective) after 3 h incubation. **(D)** Myelin phagocytosis by purified human neutrophils (n=9) over 24 h as determined through IncuCyte software. The phagocytic count was calculated considering the overlapping events between the red fluorescent myelin debris with green, fluorescent neutrophil counts. The percentage of phagocytosis was then obtained by normalizing the phagocytic count by the viable cell count. **(E-F)** Calcein-labelled purified neutrophils from healthy donors (n=9) were cultured with opsonized (serum, HI serum, PBS) pHrodo-labelled myelin debris in **(E)** buffer conditions or **(F)** in the presence of cytokines (50 ng/mL TNF-α, 10 ng/mL IFN-γ and 10 ng/mL IL-1β), whereby fluorescent images were generated every hour for 24 h with the IncuCyte system. The maximal percentage of myelin debris phagocytosis over time was determined. Normally distributed data are represented as the mean ± SEM. Data were analyzed using the paired Student’s t-test with Bonferroni correction. *p<0.05, **p<0.01, ***p<0.001. HI serum: heat-inactivated serum.

To verify the capability of human neutrophils to engulf myelin debris, an *in vitro* phagocytosis assay using pHrodo-labelled human myelin and calcein-labelled neutrophils was applied. To investigate the involvement of the complement system, myelin was opsonized with either complete human serum, containing both complement factors and immunoglobulins or heat-inactivated (HI) serum, lacking complement activation, or PBS prior to exposure to cells. Following imaging, the percentage of myelin uptake was calculated by normalizing the overlapping green (viable neutrophils) and red (internalized myelin) events by the number of viable cells at the beginning. Internalization of myelin was observed in the majority of neutrophils, which became grouped in the process (**Fig 1B**). To confirm our observations with more resolution, we imaged myelin uptake using confocal microscopy. With this method, myelin internalization by neutrophils was clearly observed, with 3D rendering confirming intracellular myelin remnants (**Fig 1C**). From 4 to 6 hours, the maximal percentage of myelin internalization was reached in all conditions (**Fig 1D**). After 7 to 9 hours, neutrophil viability started to decrease, reaching 0% survival around 19 hours, irrespective of the opsonization method (**Supplementary fig 2**). The maximal percentage of myelin uptake was found when myelin was opsonized with serum (+/- 38%) and this significantly decreased when myelin was opsonized with HI serum (+/- 15%) or PBS (+/- 18%) (**Fig 1E**). These results suggest that complement opsonizes myelin debris, driving its uptake more efficiently.

Next, to mimic the neuroinflammatory conditions observed in the brain of MS patients undergoing myelin degradation, neutrophils were activated with a mix of the MS-related cytokines TNF-α, IL-1β and IFN-γ during myelin incubation. Surprisingly, no significant difference was detected in uptake between control and cytokine-stimulated cells (**Fig 1E-F**).

Finally, we investigated whether human neutrophils also accumulate lipid droplets upon myelin exposure *in vitro* and obtain a typical foamy appearance. For this purpose, we incubated neutrophils for 4 hours with low dose (1 µg/mL) or high-dose (100 µg/mL) myelin debris and stained the cells with Nile red or Bodipy, which are lipophilic, fluorescent probes that stain neutral lipids accumulating within the core of lipid droplets (**Fig 2A-B**). We observed the formation of lipid droplets with both dyes, but only after exposure to the highest myelin dose (**Fig 2C**), confirming the ability of human neutrophils to acquire a foamy, myelin-containing phenotype when they are exposed to 100 µg/mL myelin debris.

**Figure 2.**
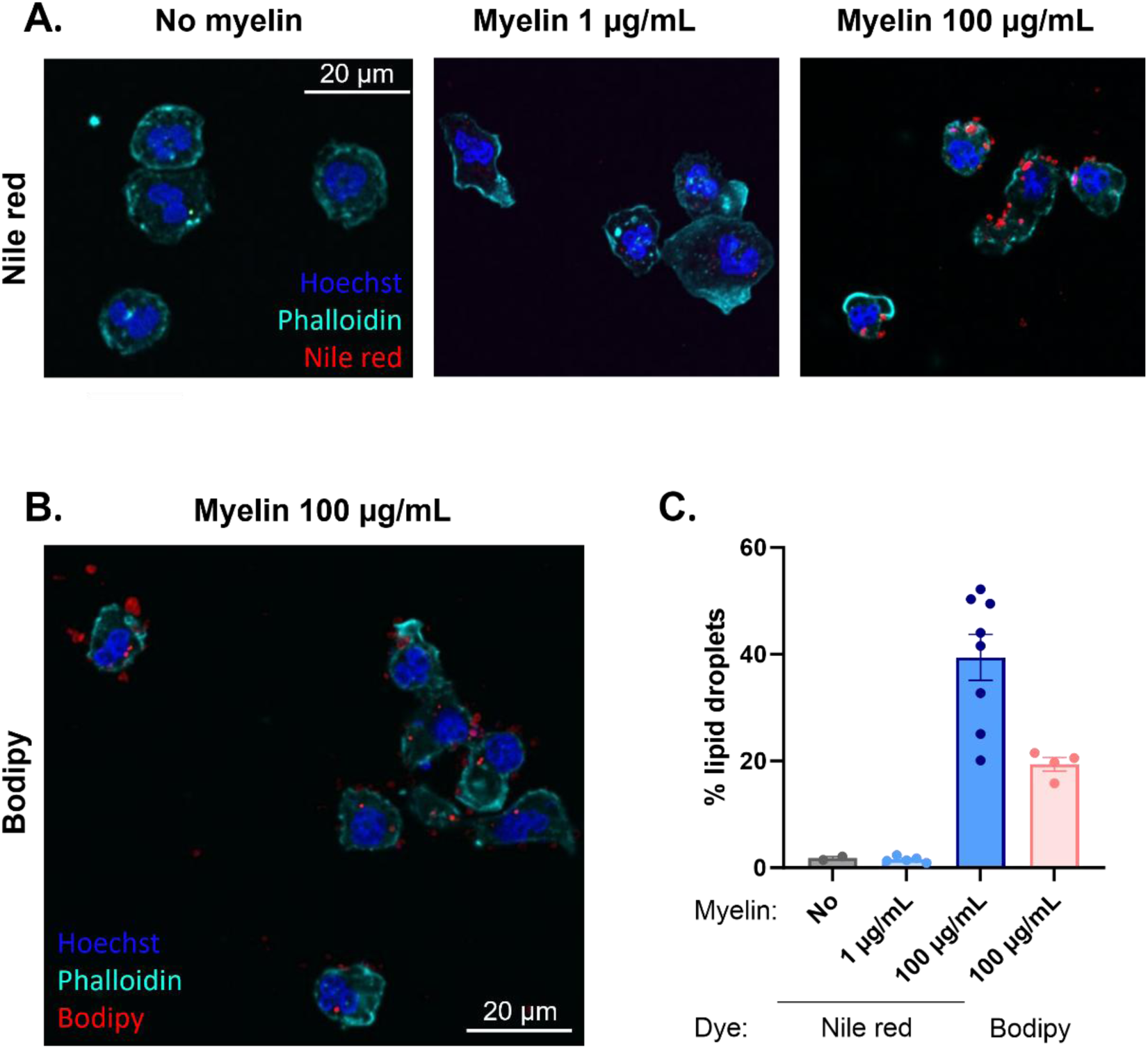
Foamy neutrophils are formed *in vitro* following high-dose myelin incubation. Human purified neutrophils from healthy donors were incubated for 4 h with unlabelled myelin debris (1 or 100 µg/mL) or PBS (no myelin condition) after which they were washed, fixed and stained with Hoechst (10 µg/mL) and Phalloidin (1:400) in combination with **(A)** Nile red (300 nM) or **(B)** Bodipy (2 µm) to stain for lipid droplets. Cells were then imaged with a confocal microscope (63x) and analyzed using ImageJ.

### 3.2 Myelin uptake by neutrophils is mediated through complement receptor 3 and soluble complement fragments

As complete human serum augmented myelin uptake by neutrophils, we investigated the possible involvement of complement receptors (CRs). Human neutrophils express various CRs, including CR3, also known as the CD11b/CD18 dimer. CR3 is a β2-integrin that is most often associated with neutrophil phagocytosis, as it recognizes iC3b-opsonized pathogens, particles and cell debris (26). To determine the importance of CR3 in driving myelin uptake by neutrophils, cells were pre-treated with an anti-CD11b blocking antibody. The percentage myelin uptake in untreated cells (PBS) or the isotype-treated group reached a maximum of 30-40% (**Fig 3A**). Blocking CR3 significantly decreased myelin uptake compared to the serum-PBS and serum-isotype controls, reducing it to levels observed with non-opsonized myelin. Since the ligand for CR3 is the activated complement fragment iC3b and, to a lesser extent C1q, we repeated the phagocytosis assay, opsonizing the myelin debris with either complete human serum or C3-/C1q-depleted serum. As a control, the cytoskeleton inhibitor latrunculin B was included, which prevents polymerization of actin filaments and thereby engulfment of particles. As previously observed, the maximal percentage internalization of myelin opsonized with normal serum peaked at 35%, which was significantly reduced when myelin was opsonized with C3- or C1q-depleted serum, or latrunculin B (**Fig 3B**). To investigate the involvement of Fc receptors in myelin internalization, neutrophils were also pre-treated with excess IgG to block Fcγ receptors, however no effect on myelin uptake was observed (**Supplementary fig 3**).

**Figure 3.**
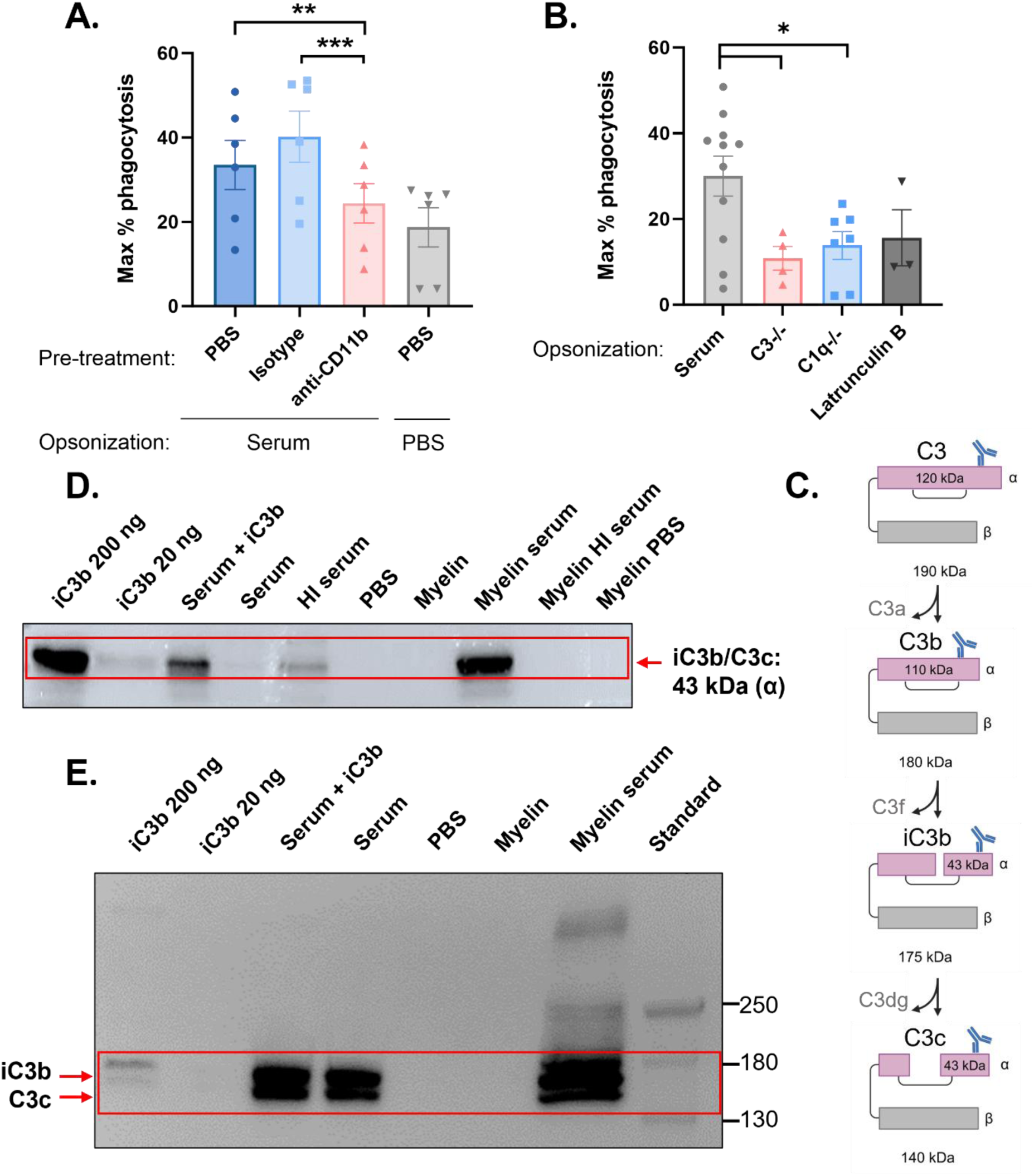
Phagocytosis of myelin debris by neutrophils involves complement receptor 3 and C3/C1 fragments. **(A)** Calcein-labelled purified neutrophils from healthy donors (n=6) were pre-treated with anti-CD11b (CR3) antibody, IgG2b isotype or PBS control, for 15 min and cultured with PBS- or serum-opsonized pHrodo-labelled myelin debris (75 µg/mL). Fluorescent images were generated every hour for 24 h with the IncuCyte imaging system. Thereafter, maximal percentage of myelin debris phagocytosis was determined. **(B)** Calcein-labelled purified neutrophils from healthy donors (n=3-7) were pre-treated for 15 min with latrunculin B or PBS and afterwards incubated with pHrodo-labelled myelin debris (75 µg/mL), opsonized with serum, C3-depleted or C1q-depleted serum. Maximal percentage of myelin debris phagocytosis is shown. Normally distributed data are represented as the mean ± SEM. Data were analyzed using the paired Student’s t-test with Bonferroni correction. *p<0.05, **p<0.01, ***p<0.001. **(C)** Structural composition of complete C3 (190 kDa) and its subsequent cleavage to fragments C3b (180 kDa), iC3b (175 kDa) and C3c (140 kDa). The image shows the localization of the binding epitope for the C3 antibody used in this study, at the C-terminal side (360 AA) of the common α chain in all fragments. **(D,E)** Western blot analysis of human myelin in **(D)** reducing and **(E)** non-reducing conditions. The purified recombinant human iC3b also contains some C3c peptide. This preparation and serum (1/200 diluted) enriched with recombinant iC3b were included as positive controls. HI serum: heat-inactivated serum.

Having established that the complement protein C3 within serum and the complement receptor CR3 on neutrophils are important for myelin uptake by neutrophils, we next assessed the mechanism of interaction between myelin and the components of the complement system. The full C3 fragment (190 kDa) can be sequentially cleaved inside its α chain to form multiple smaller C3-derived peptides (**Fig 3C**) (27). Myelin was opsonized with serum, HI serum or PBS and washed extensively afterwards, to remove unbound particles. Next, the mixtures were separated with gel electrophoresis in a reducing or non-reducing environment, after which the samples were blotted onto a membrane and stained with an anti-C3b antibody. This antibody recognizes the α chain of C3 and its cleavage products C3b, iC3b and C3c (**Fig 3C**). First, a reducing loading buffer was used, which disrupts disulfide bridges between the α and β chain of all C3 fragments and all non-covalent protein-protein interactions (**Fig 3D**). A mixture of recombinant iC3b and C3c (labelled as iC3b in Fig 3) was used as positive control and was also spiked in human serum. A band corresponding to the α chain of iC3b (∼43 kDa) is visible in the lane of the positive control and the serum spiked with iC3b/C3c. As expected, no signal was observed in the negative control (PBS lane). In the myelin lanes, iC3b signal is only visible in the serum-opsonized myelin condition and absent in pure myelin or myelin opsonized with HI serum or PBS. In reducing conditions, we cannot distinguish iC3b from C3c, as the antibody recognizes an epitope in the α chain which is identical for both. Therefore, a non-reducing loading buffer was used, keeping disulfide bridges and non-covalent interactions intact (**Fig 3E**). This time, the positive control showed two bands, around 176 kDa and 140 kDa, corresponding to iC3b and C3c, respectively. The same bands are visible in the serum and the spiked (iC3b/C3c) serum lanes but absent in the PBS or pure myelin lanes. In the serum-opsonized myelin condition, two bands were detected with an apparent molecular size identical to the lanes containing the pure iC3b/C3c and enriched serum. However, a highly pronounced third band (∼180 kDa) and a faint fourth band (∼ 250 kDa) can be observed in the serum-opsonized myelin lane, which are not present in any of the other lanes. These bands could represent C3 fragments bound to myelin or remaining intact C3 protein (∼ 190 kDa). These experiments confirm that active cleavage products of C3 (such as iC3b) opsonize myelin covalently.

### 3.3 Myelin internalization induces a pro-inflammatory gene signature in foamy neutrophils

To reveal whether myelin uptake affects the transcriptome of neutrophils, purified neutrophils from five different healthy donors were exposed to PBS or two doses of myelin (1 or 100 µg/mL). After 12 h of myelin exposure, mRNA was isolated and bulk RNA sequencing was performed. The resulting heat map of the 500 genes with highest variance across different groups showed that incubation with low-dose myelin (1 µg/mL) did not induce major changes in the neutrophils’ transcriptome compared to PBS-control cells (**Fig 4A**). In contrast, high-dose myelin (100 µg/mL) caused a complete shift in gene expression profile. When focusing on expression of inflammatory genes (**Fig 4B**), the immunoregulatory gene *ARG1* was amongst the top down-regulated genes upon high-dose myelin (100 µg/mL) stimulation. Furthermore, both *CASP1* and *FAS* showed decreased expression in the high-dose myelin group, indicating resistance to cell death. In contrast, several inflammatory cytokines (*CCL3*, *IL-1B*, *CCL4*, *CCL4L2*), molecules and enzymes (*S100A11, S100A9, PADI4, MMP9, MPO*), were upregulated. Pathway analysis on the top 100 differentially expressed genes (high-dose myelin vs no myelin) revealed activation of pathways involved in fatty acid oxidation, cholesterol metabolism and lipid transport (**Fig 4C**). This matches our previous observations that myelin engulfment impacts the metabolic phenotype of myelin-induced foamy microglia (28).

**Figure 4.**
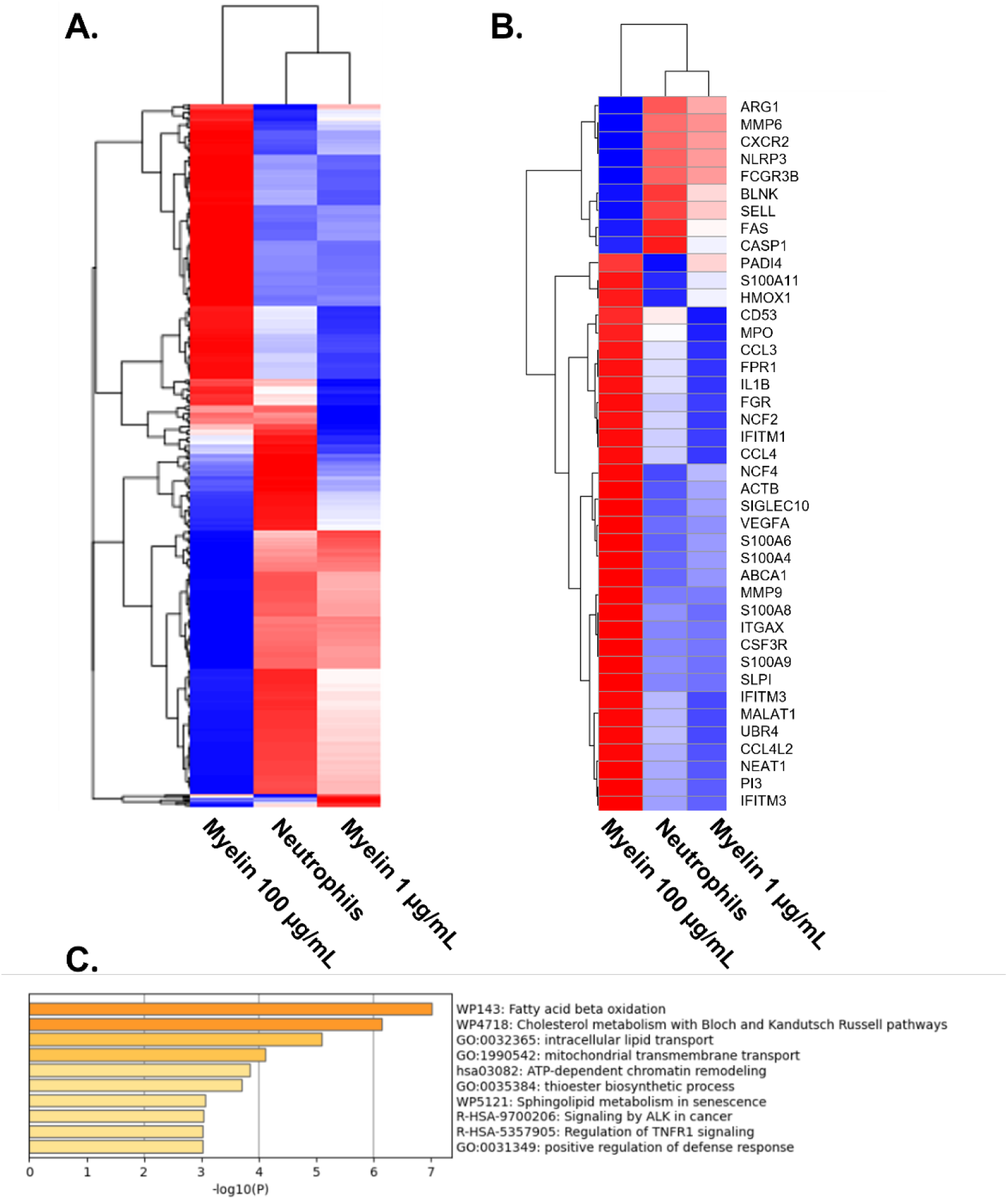
Bulk RNA sequencing reveals an inflammatory gene expression profile in neutrophils subject to high-dose myelin uptake. Purified neutrophils from healthy donors (n=5) were incubated for 12 h with PBS or myelin debris (1 or 100 µg/mL) after which RNA was isolated and sequenced. **(A)** Heatmap from the top 500 genes with highest variance amongst all groups. **(B)** Heatmap showing pre-selected inflammation-related genes. **(C)** Pathway analysis on the top 100 differentially expressed genes between the high-dose myelin (100 µg/mL) and neutrophils-only group.

To further explore the gene signature identified by RNA sequencing, we performed qPCR analysis of key pro- and anti-inflammatory markers following exposure to different myelin doses. As a separate validation step, we first confirmed that these genes respond to inflammatory stimulation at the selected timepoints. To this end, neutrophils were stimulated for 6 or 12 h with an inflammatory cytokine mix consisting of TNF-α, IFN-γ and IL-1β. This treatment induced the expected upregulation of *CXCL8, HMOX1, TNF* and *VEGF* mRNA expression, which was most pronounced at 12 h (**Supplementary fig 4**). In contrast, the anti-inflammatory genes *ARG1* and *TGFB1* were significantly downregulated upon cytokine stimulation, although the effect on *TGFB1* was no longer evident at 12 h (**Supplementary fig 4**). Having established the responsiveness of these markers, we next assessed their expression following myelin uptake. Neutrophils were incubated with PBS or increasing doses of serum-opsonized myelin debris (1, 10 or 100 µg/mL). After 12 h, stimulation with the highest myelin concentration (100 µg/mL) resulted in a significant upregulation of *CXCL8* and *HMOX1,* along with a trend toward increased *TNF* and *VEGF* expression (**Fig 5**). In contrast, *ARG1* expression was significantly decreased following exposure to the highest myelin dose at both 6 and 12 h (**Fig 5**).

**Figure 5.**
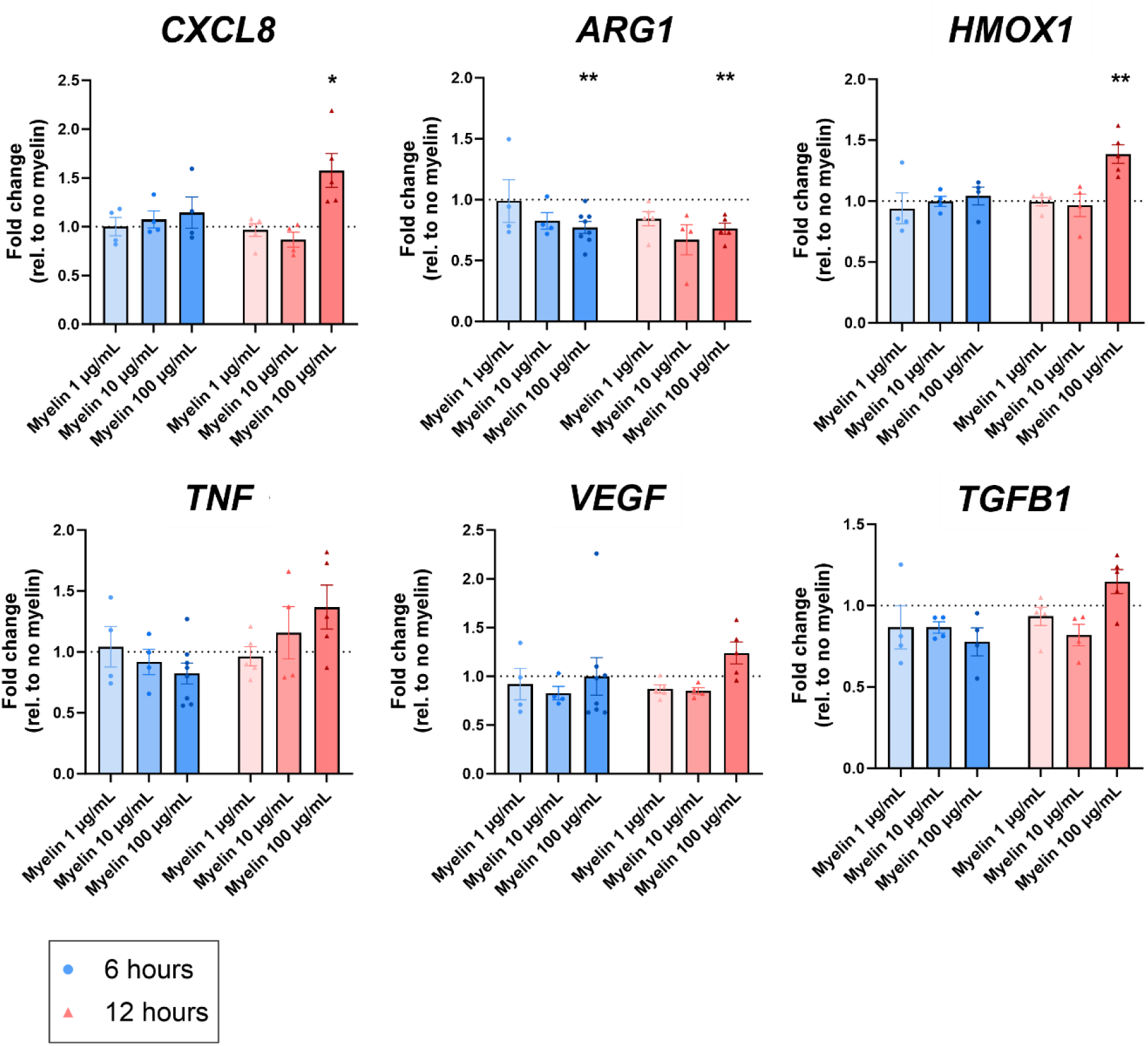
Pro-inflammatory versus pro-resolving gene expression in human neutrophils after phagocytosis of myelin. Human neutrophils (n=4-8) were incubated for 6 or 12 h with different doses of serum-opsonized myelin debris (1, 10 or 100 µg/mL). Afterwards, mRNA was isolated, cDNA was produced, and qPCR was performed. Data were normalized to the expression of the housekeeping gene 18S and are represented as fold change relative to buffer-stimulated neutrophils and represented as the mean ± SEM. Normally distributed data were analyzed using the one-sample Student’s t-test, with comparison value set at 1; *p<0.05, **p<0.01.

### 3.4 The release of chemokines and proteases is increased by myelin uptake in a dose-dependent manner

To further investigate the effect of myelin uptake on the inflammatory properties of neutrophils and confirm the RNA sequencing data, the levels of cytokines and proteases were quantified using ELISA in the supernatants of neutrophils that were stimulated for 2 h (proteases) or 24 h (cytokines) with different doses of myelin debris (1, 10 or 100 µg/mL). The levels of IL-10, CXCL10 and TGF-β were below the detection limit. The concentrations of the pro-inflammatory chemokines CXCL8 and CCL3 were significantly increased after stimulation of neutrophils with myelin debris, in a dose-dependent manner (**Fig 6A,B**). The highest levels of chemokines were always measured in supernatants of neutrophils that were challenged with 100 µg/mL myelin. Similarly, the level of neutrophil elastase (NE) and the activity of MMP-9 in the supernatants of neutrophils were significantly higher when cells were incubated with 100 µg/mL myelin (**Fig 6C,D**). Collectively, these data indicate that myelin uptake drives a dose-dependent increase in pro-inflammatory chemokine release and protease activity in neutrophils.

**Figure 6.**
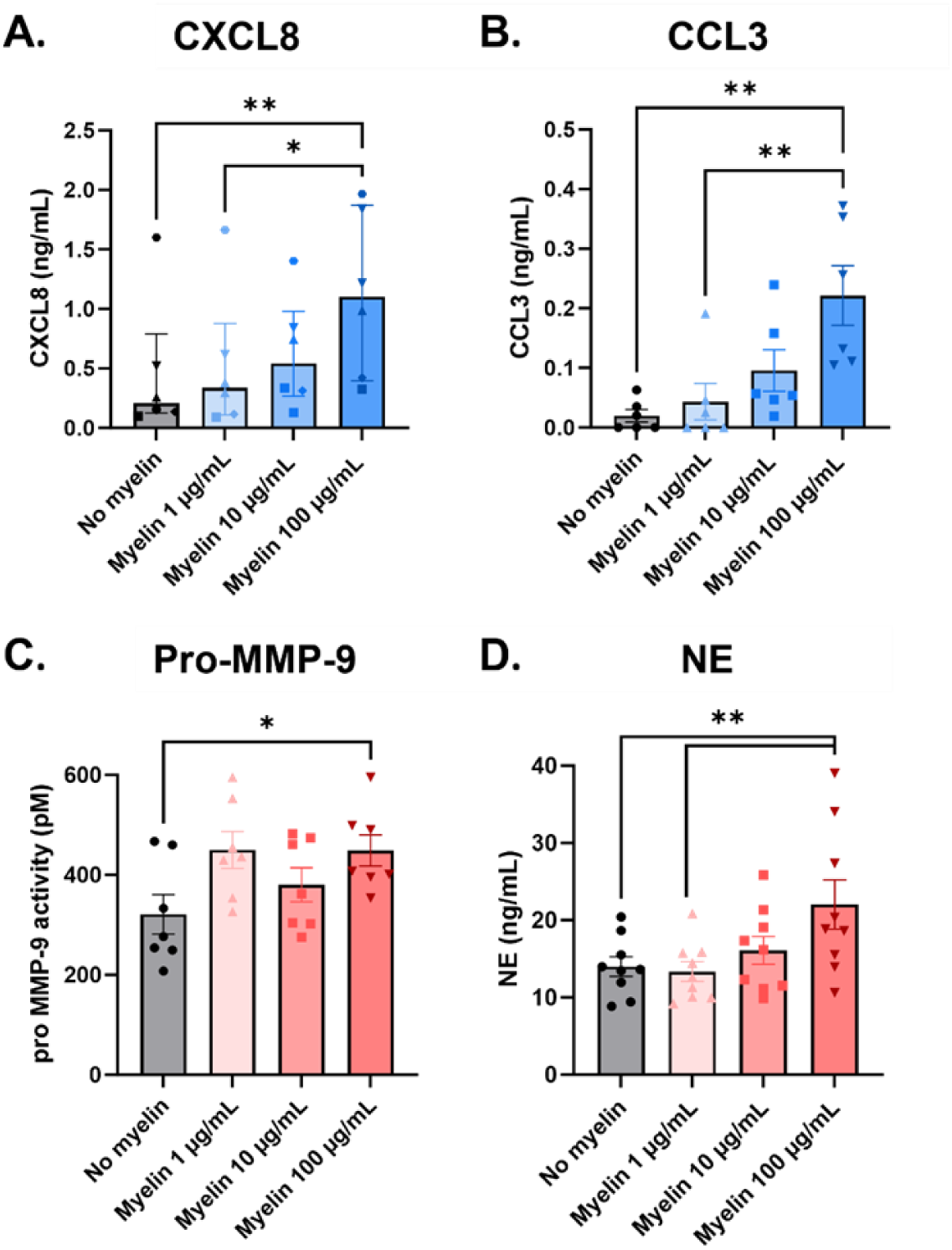
The production of CXCL8, CCL3, pro-MMP-9 and neutrophil elastase (NE) increases in a dose-dependent manner after myelin uptake by neutrophils. Purified neutrophils from healthy donors (n=6-9) were cultured with or without serum-opsonized myelin debris at different concentrations (1, 10 or 100 µg/mL) for 2 h (pro-MMP-9 and NE) or 24 h (CXCL8 and CCL3). Afterwards, the concentration of CXCL8, CCL3 or NE in the supernatant was determined with ELISA. The enzymatic activity of pro-MMP-9 in the supernatant was quantified with gelatin zymography. Normally distributed data are represented as the mean ± SEM and non-normally distributed data as median ± interquartile range. Data were analyzed using a One-way ANOVA with Dunnett’s correction (normally distributed data) or the Friedman test with Dunn’s correction (non-normally distributed data). *p<0.05, **p<0.01

### 3.5 Myelin uptake drives the production of reactive oxygen species (ROS) and neutrophil extracellular traps (NETs)

Next, we measured ROS production after short (6 h) and long (24 h) exposures of neutrophils to different concentrations of myelin debris (1, 10 or 100 µg/mL). Following incubation for the respective time periods, neutrophils were harvested and stimulated with buffer, PMA or cytokines (10 min TNF-α priming + IL-1β stimulation) after which ROS production was quantified using chemiluminescence. At the 6 h timepoint, no significant differences could be observed between the different myelin doses, irrespective of the applied stimulus (**Supplementary fig 5**). After 24 h, spontaneous ROS production in the buffer group was low, with a clear increase in signal upon stimulation with cytokines or PMA in the absence of myelin. Additionally, neutrophil ROS production was significantly elevated upon stimulation with myelin in a dose-dependent manner, compared to untreated cells or lower myelin concentrations (**Fig 7A**). ROS production was further increased in response to PMA or TNF-α + IL-1β also dose-dependently, with the highest signal obtained in the 100 µg/mL myelin treatment groups.

**Figure 7.**
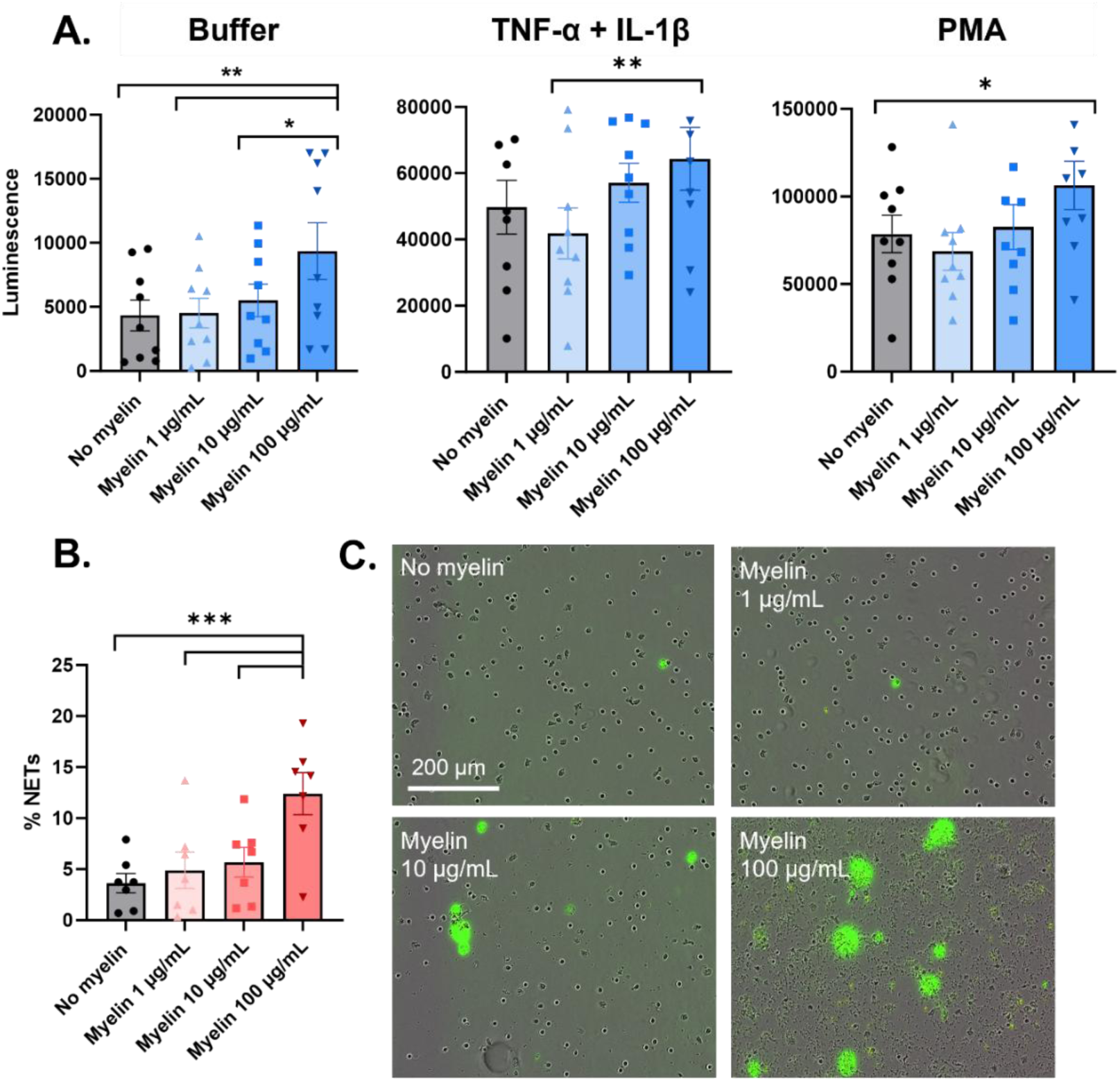
The production of reactive oxygen species and neutrophil extracellular traps increases in a dose-dependent manner after myelin uptake. **(A)** Purified neutrophils from healthy donors (n=5-6) were cultured with or without serum-opsonized myelin debris at different concentrations (1, 10 or 100 µg/mL) for 24 h. Afterwards, cells were harvested, counted and a chemiluminescence-based ROS assay was performed. Therefore, 0.1 x 10^6^ cells were added to each well, and stimulated with RPMI buffer, PMA (150 ng/mL) or a combination of cytokines (priming for 10 min with 10 ng/mL TNF-α and subsequent stimulation with 10 ng/mL IL-1β). Afterwards, luminol (2 mM) was added to the cells, and luminescence was measured for 3 h at 37°C in a CLARIOstar microplate reader. Maximal luminescence values over time are shown. Normally distributed data are represented as the mean ± SEM. Data were analyzed using a one-way ANOVA. *p<0.05, **p<0.01. **(B)** Purified neutrophils from healthy donors (n=7) were added to a 96-well plate (5 x 10^4^ cells/well) and stained with SYTOX green (50 nM) for 30 min at 37°C. Afterwards, cells were stimulated with different doses of serum-opsonized myelin debris (1, 10 or 100 µg/mL) and this was imaged each hour for 10 h at 37°C using the IncuCyte system. The percentage NETs release over time was calculated by dividing the NETs area (SYTOX green) over the cellular area at timepoint 0 h. The increase in percentage NETs release (T=10h – T=0h) is shown. Normally distributed data are represented as the mean ± SEM. Data were analyzed using a one-way ANOVA. ***p<0.001. **(C)** Representative IncuCyte images after 3 h of phagocytosis are shown, where extracellular released DNA during NETosis is stained using SYTOX green, with 20x magnification.

As ROS production is one of the crucial steps in the formation of NETs, and NETs are found to be elevated in the circulation of MS patients (13), we decided to evaluate whether myelin uptake promotes the production of NETs. The release of NETs in response to increasing serum-opsonized myelin concentrations (1, 10 and 100 µg/mL) was visualized using SYTOX green, and quantified with the IncuCyte imaging system (**Fig 7B,C**). Our results show a dose-dependent increase in NET release, which was significantly higher when the highest concentration of myelin was added to neutrophils (**Fig 7B,C**). Overall, internalization of myelin stimulates major effector mechanisms in neutrophils, ROS production and NETosis, both of which are contributors to tissue damage.

## 4 Discussion

Neutrophils, as professional phagocytes of the innate immune system, are often neglected in MS research, despite their presence in the CNS during the early and peak stages of disease in both EAE and cuprizone mouse models. In this study, we explored the potential role of human neutrophils in the clearance of myelin debris. We examined human and mouse lesions to demonstrate neutrophil involvement. Additionally, we examined whether the uptake of myelin induced changes in the phenotype and functions of neutrophils and revealed which receptors and pathways are involved. Investigating the dynamics of myelin uptake in MS context improves our understanding of the balance between myelin debris clearance, supporting remyelination *versus* possible pro-inflammatory disease promoting myelin uptake (7,29).

Confirming literature (30), we observed neutrophil infiltration in the inflamed spinal cord tissue of EAE mice, mainly in regions characterized by active demyelination. In this model, Yamasaki *et al.* detected murine neutrophils engulfing myelin with electron microscopy (10). Additionally, Lindborg *et al.* demonstrated that neutrophils are essential for the clearance of inhibitory myelin debris in Wallerian degeneration in the peripheral nervous system (31). In support of this notion, we demonstrate the presence of “foamy” lipid-droplet-associated neutrophils in demyelinated regions of human postmortem MS lesions. We confirmed this phenomenon *in vitro*, where neutrophils exposed to myelin debris accumulated abundant lipid droplets. When studying kinetics of myelin uptake, we found that human neutrophils take up myelin debris, reaching a peak after 3 to 6 hours. Previous in vitro studies have shown that myelin debris is also efficiently taken up by other phagocytes, such as microglia and macrophages, with uptake capacity varying by cell type (32). In the EAE mouse model for MS, myelin uptake was observed starting at day 5 in the spinal cord tissue, with microglia dominating uptake during the initial phase (33). Hendrickx et al. also used a pHrodo-based myelin phagocytosis assay, showing that THP-1-derived macrophages exhibit rapid myelin internalization, reaching maximal phagocytosis after 24 hours. On the other hand, primary human microglia display a slower but sustained uptake of myelin for up to 70 hours (34). In our experiments, neutrophils were also capable of substantial myelin phagocytosis, which depended on myelin dose and serum opsonization. Notably, phagocytic capacity declined after several hours, likely reflecting the limited lifespan of neutrophils.

Through the comparison of different opsonization methods, we demonstrated that opsonization of myelin debris with complete human serum augmented phagocytosis (30-40%), which was significantly higher compared to HI serum or PBS (10-15%). These results are in line with data from macrophage and microglia research, showing that the presence of intact complement enhances phagocytosis of myelin *in vitro* (35,36). Blocking CR3 (CD11b/CD18) reduced myelin phagocytosis, further confirming complement involvement, identifying its ligand iC3b as possible active opsonin in this process. Notably, the maximal percentage of phagocytosis when myelin was opsonized with C1q- and C3-depleted serum was lower compared to the cells treated with anti-CR3, indicating that the complement proteins C1q and C3 are necessary for myelin uptake, but additional complement receptors binding these factors (e.g., CR4 and to a lesser extent CR1) may also be involved. These data are consistent with findings in macrophages and microglia, where blockage of CR3 also resulted in inhibition of phagocytosis (35,37,38). Further analysis of the possible interactions between C3 fragments and myelin using immunoblot confirmed binding of iC3b to myelin, which is then recognized by CR3 for engulfment. The importance of C3 for myelin uptake *in vivo* was previously reported as C3 knockout mice or the use of pharmacological C3 inhibitors reduced myelin phagocytosis by macrophages and protected mice from EAE disease development, synapse loss and microglia activation (18,39). Moreover, complement deposition was detected in demyelinating lesions from MS patients and complement upregulation by neuronal cells has also been implicated in demyelinating disorders. Furthermore, in MS patients, C3 levels are elevated in CSF and active demyelinating lesions are positive for C3, C3b, iC3b and C4b (40,41). No effect was observed on myelin phagocytosis after blocking Fc receptors with an excess IgG (**Supplementary fig 3**), excluding Fc related pathways for myelin uptake in neutrophils. Of interest, other studies showed that anti-myelin specific antibodies boosted phagocytosis in macrophages (36,42).

As we could still detect some remaining internalization when myelin was opsonized with HI serum, C3- or C1q-depleted serum, we cannot rule out that other receptors besides complement receptors are involved. For instance, scavenger receptors (SR) such as SR-AI/II, CD36, LDL receptor related protein 1 (LRP1), collectin placenta 1 and lectin-type oxidized LDL receptor 1 (LOX-1) were demonstrated to be involved in myelin uptake by macrophages (16). However, while SR-AI/II are highly expressed on foamy macrophages and ramified microglia around chronic active MS lesions (43), their expression on neutrophils remains debatable (44,45). Scavenger receptor class B, CD36, mainly recognizes oxidized LDL and blocking this receptor resulted in increased neuroinflammation, both *in vitro* and in the EAE model (46). This fatty acid translocase is expressed at low levels on resting neutrophils but was found to be upregulated upon activation and on low-density neutrophils (47). For instance, after intracellular accumulation of cholesterol or following cytokine activation, neutrophils upregulated CD36 and LRP1 (48,49). Proof of the involvement of LOX-1 in myelin phagocytosis remains lacking. However, its expression is upregulated at sites of active demyelination in MS lesions (43). LOX-1 expression is elevated on activated neutrophils in diseases such as COVID-19, cancer and sepsis (50). In mice, *Lox-1* deletion improved neutrophil recruitment and resolution of inflammation in pneumonia models, and LOX-1^+^ neutrophils upregulated a gene signature related to cholesterol metabolism (51,52).

Next, we wanted to explore the possibility of a phenotype switch in neutrophils induced by myelin phagocytosis as seen in macrophages and microglia, investigating different timepoints and concentrations. As ROS production has been directly linked to myelin phagocytosis (53,54), and neutrophil-derived ROS damages the BBB in the EAE model (12), we first investigated ROS production following myelin internalization. Long-term (24 h) exposure of myelin to neutrophils, increased both spontaneous and PMA- or cytokine-induced ROS release. This is in line with data on microglia, showing a dose-dependent increase in ROS production following myelin uptake (53). The increase in oxygen reactants after myelin internalization might be mediated by CR3, and subsequent activation of the NADPH oxidase pathway (55,56). Interestingly, ROS production may also directly damage the myelin sheath through the oxidation of lipids, thereby promoting further breakdown of myelin and inflammation in the CNS (57). This was evidenced in MS patients, where lipid peroxidation was detected in the CSF (58). Also, in MS brain tissue, demyelination and neurodegeneration were found to be closely associated with oxidized lipids in myelin membranes and oxidized LDL was found in actively demyelinating plaques as well as in foamy macrophages (59,60). Lastly, ROS production is directly toxic for oligodendrocytes, causing cellular apoptosis in both human primary oligodendrocytes and in the cuprizone mouse model for demyelination (61,62).

As ROS production precludes the formation of NETs, which are also linked to autoimmunity, we investigated NET release by neutrophils phagocytosing myelin. NETosis is normally induced in response to microbial patterns or endogenous danger motifs, and its release must be tightly balanced, as uncontrolled NET release causes tissue damage, for example during autoimmunity or chronic inflammation (63). NETs are formed after activation of surface receptors on neutrophils, leading to an intracellular cascade that includes ROS production and the release of myeloperoxidase (MPO), NE and protein arginine deiminase 4 (PAD4), the latter of which is also known to citrullinate MBP in MS patients (64,65). In line with the data on ROS production, the release of NETs significantly increased when neutrophils were cultured in the presence of 100 µg/mL myelin. This highlights again the pro-inflammatory and potentially damaging phenotype of neutrophils stimulated with high-dose myelin. Notably the *PADI4* gene, encoding the PAD4 enzyme which citrullinates histones during NETosis, was highly upregulated in neutrophils challenged with high doses of myelin.

Prolonged exposure of neutrophils to high myelin concentrations induced a pro-inflammatory phenotype, as confirmed by gene expression analysis. Upregulation of the pro-inflammatory chemokine CXCL8 can amplify the attraction of additional neutrophils and thereby maintain a pro-inflammatory environment (66,67). While HMOX1 is often considered anti-inflammatory in macrophages, in neutrophils it likely reflects a stress-responsive role linked to oxidative and inflammatory signaling. It was significantly upregulated after long exposure with the highest myelin concentration. Interestingly, genetic variants of *HMOX1* were also identified as significant risk factors for MS (68). In contrast, the anti-inflammatory marker *ARG1* was consistently downregulated, supporting a shift toward a pro-inflammatory (N1-like) phenotype following myelin uptake. (69,70).

These transcriptional changes were mirrored at the protein level. Myelin uptake induced a dose-dependent increase in CXCL8, CCL3, neutrophil elastase (NE), and MMP-9 release, with the strongest effects at the highest myelin concentration. CXCL8 and CCL3 are well-established inflammatory mediators, previously shown to be elevated in the CSF, serum and brain tissue of MS patients (71–74). This increased chemokine release suggests that myelin-exposed neutrophils can amplify inflammation. CCL3, in particular, is implicated in myeloid cell activation and has been linked to demyelination in experimental models (75–77). Increased MMP-9 and NE further point to enhanced tissue remodeling capacity, consistent with their known association with BBB disruption and disease activity in MS (13,78–80).

Altogether, these observations demonstrate that neutrophils switch towards a proinflammatory phenotype upon myelin debris uptake in a dose- and time-dependent way. This is in line with macrophages and microglia, which are skewed to a more pro-resolving, beneficial phenotype after initial myelin internalization (up to 24 hours). However, long-term (> 72 hours) myelin uptake eventually pushes these cells to an M1 pro-inflammatory, repair-inhibitory phenotype, in a dose-dependent manner. This is caused by dysregulation of metabolic pathways involved in myelin processing and efflux, ultimately resulting in the formation of foamy macrophages/microglia (7,29,81).

## Conclusion

This study demonstrates that foamy, lipid-loaded neutrophils develop within demyelinated regions of CNS lesions in both EAE and MS tissue. Further, human neutrophils can phagocytose myelin debris, a process partially mediated by iC3b binding to CR3. Myelin internalization induced a pro-inflammatory phenotype in neutrophils in a dose- and time-dependent manner, with the most pronounced effects observed after 24 hours of incubation with myelin debris. We detected a dose-dependent increase in both spontaneous-and stimulus-induced (TNF-α + IL-1β or PMA) ROS production in neutrophils incubated with myelin compared to unstimulated cells. Moreover, 100 µg/mL myelin triggered enhanced NET release. Additionally, myelin exposure enhanced the production of inflammatory mediators (chemokines and proteases) in a dose-dependent manner. These data, together with the inflammatory gene signature observed upon neutrophil exposure to 100 µg/mL myelin indicate that myelin debris drives a shift towards a pro-inflammatory and activated phenotype in neutrophils, with the strongest response occurring after prolonged incubation with high myelin concentrations. These results provide new insights into the mechanisms underlying initial myelin breakdown and the subsequent pro-inflammatory responses observed in MS and its animal models.

## Acknowledgements

The authors thank all the healthy volunteers who donated blood for this research and Dr. Gaël Vermeersch, Helga Ceunen, Anneleen Gerits and Shannon Nicolai for assistance in collecting blood samples. We thank Prof. Laurent Gillet for the mouse tissue samples and Katrien Wauterickx for performing microscopy staining. We also want to acknowledge the efforts of Lena De Waele for invaluable technical support. Lastly, we wish to thank Prof. Patrick Matthys for his valuable support and opinion on the experimental set-up and data. This work was financially supported by grants from the Research Foundation of Flanders (G025923N), Bijzonder Onderzoeksfonds, UHasselt, KU Leuven, and the Belgian Charcot Foundation. MDB and JR were financially supported by a personal grant from the Research Foundation of Flanders (1192221N and 1134425N).

## Supplementary methods

### Immunofluorescence staining of spinal cord tissue from experimental autoimmune encephalomyelitis (EAE) mice

Mice were sacrificed at day 15 post EAE induction and spinal cord tissue was snap frozen. Sections were prepared and dried at RT for 30 min, after which they were fixed in cold acetone (4°C) for 10 min, and subsequently dried at RT for 30 min. Sections were washed three times for 5 min with PBS and blocked with 0.05% Triton-X and 10% Dako protein block (Agilent) in PBS for 20 min at RT. Sections were incubated overnight at 4°C with following primary antibodies: rat anti-MBP (Sigma-Aldrich #MAB386; 1/500 in block buffer) and rabbit anti-Ly6G (Fisher scientific, Hampton, NH, USA; Clone 1A8, 1/500 in block buffer), followed by 3 PBS washes. Sections were then incubated with following secondary antibodies: AF555 goat anti-rat (1/400 in PBS, Invitrogen) and AF488 goat anti-rabbit (1/500 in PBS, Invitrogen) for 1 h at RT, washed and stained with DAPI for 10 min at RT. Finally, sections were incubated with 0.3% Sudan Black (Merck, Darmstadt, Germany) in 70% ethanol to limit autofluorescence and mounted with Fluoromount-G™ Mounting Medium (Invitrogen). Sections were imaged using a Leica DM2000 LED microscope (Leica Microsystems, Wetzlar, Germany).

## Supplementary figures

**Supplementary Figure 1.**
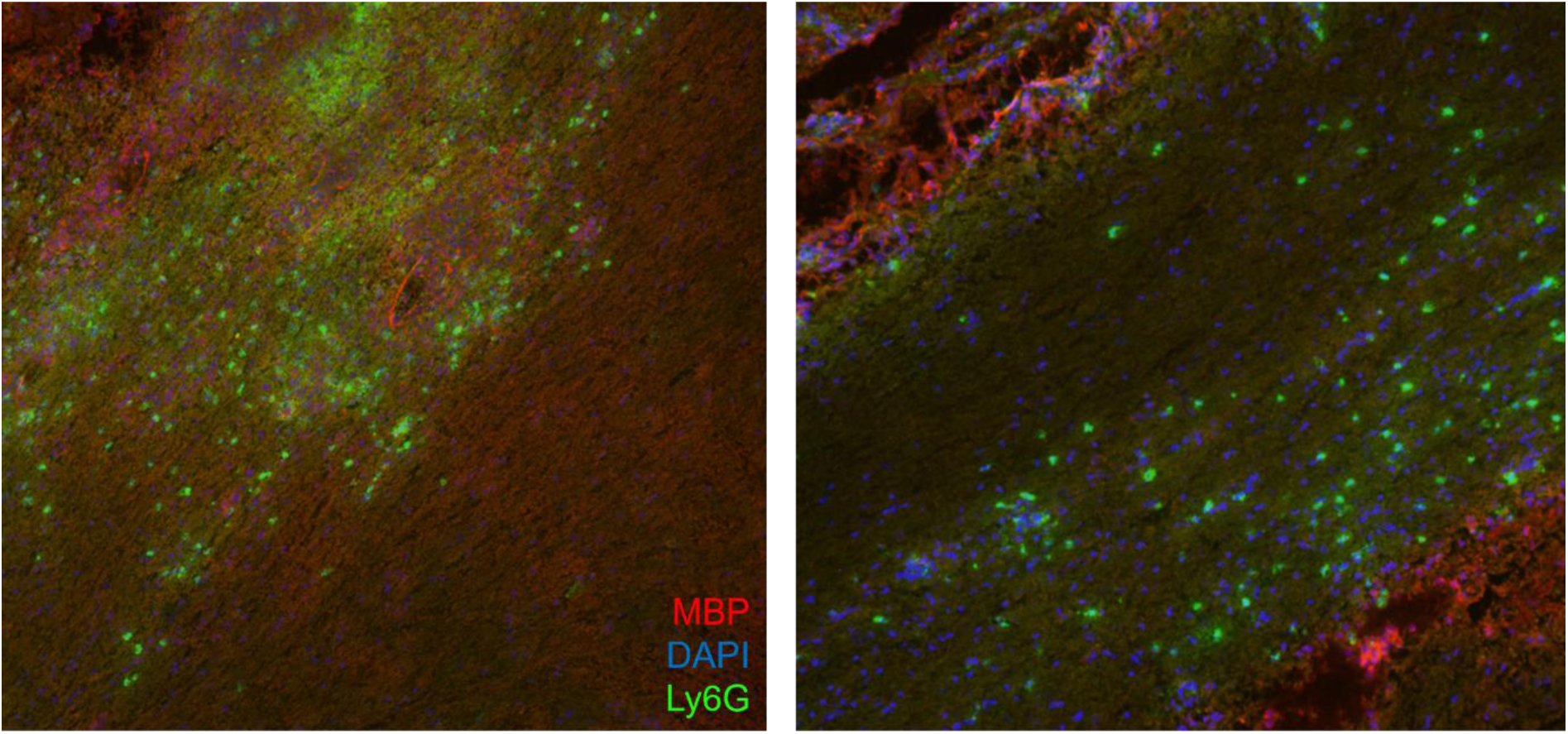
Neutrophils are present in EAE spinal cord tissue in demyelinated regions. Mice with EAE (n=6 per group) were sacrificed at day 15 post EAE induction and spinal cord tissue was fixed (10% paraformaldehyde) and stained. Red = MBP (555), Blue = DAPI, Green = Ly6G (488)

**Supplementary figure 2.**
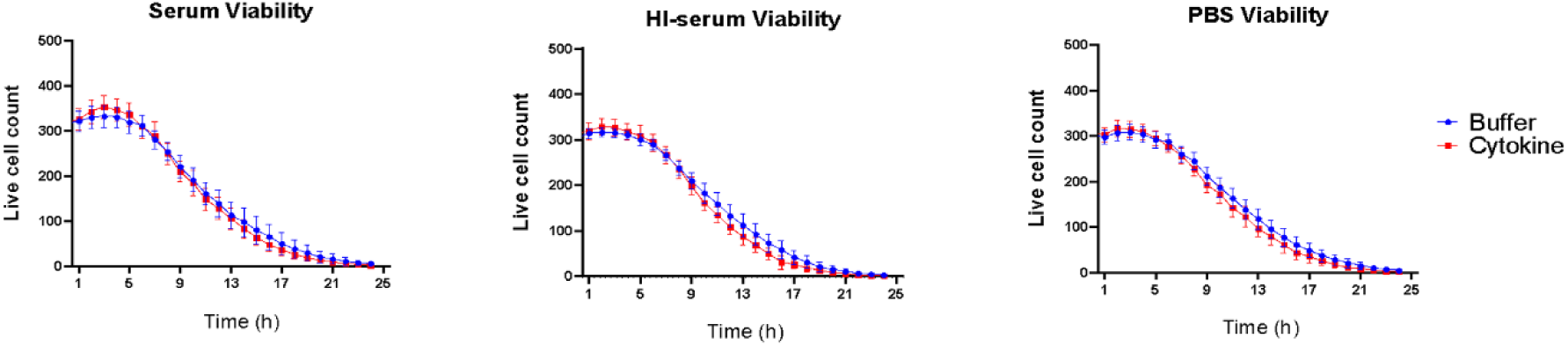
Effect of opsonized myelin on neutrophil viability over time. Calcein-labelled purified neutrophils (n=5) from healthy donors were incubated with pHrodo-labelled myelin debris, whereby fluorescent images were generated every hour for 24 h with the IncuCyte imaging system. Viability of human neutrophils (calcein green positive cells) in buffer conditions (PBS) and after addition of cytokines (50 ng/mL TNF-α, 10 ng/mL IFN-γ and 10 ng/mL IL-1β) over 24 h, analyzed with the IncuCyte imaging system. Normally distributed data are represented as the mean ± SEM. HI serum: heat-inactivated.

**Supplementary figure 3.**
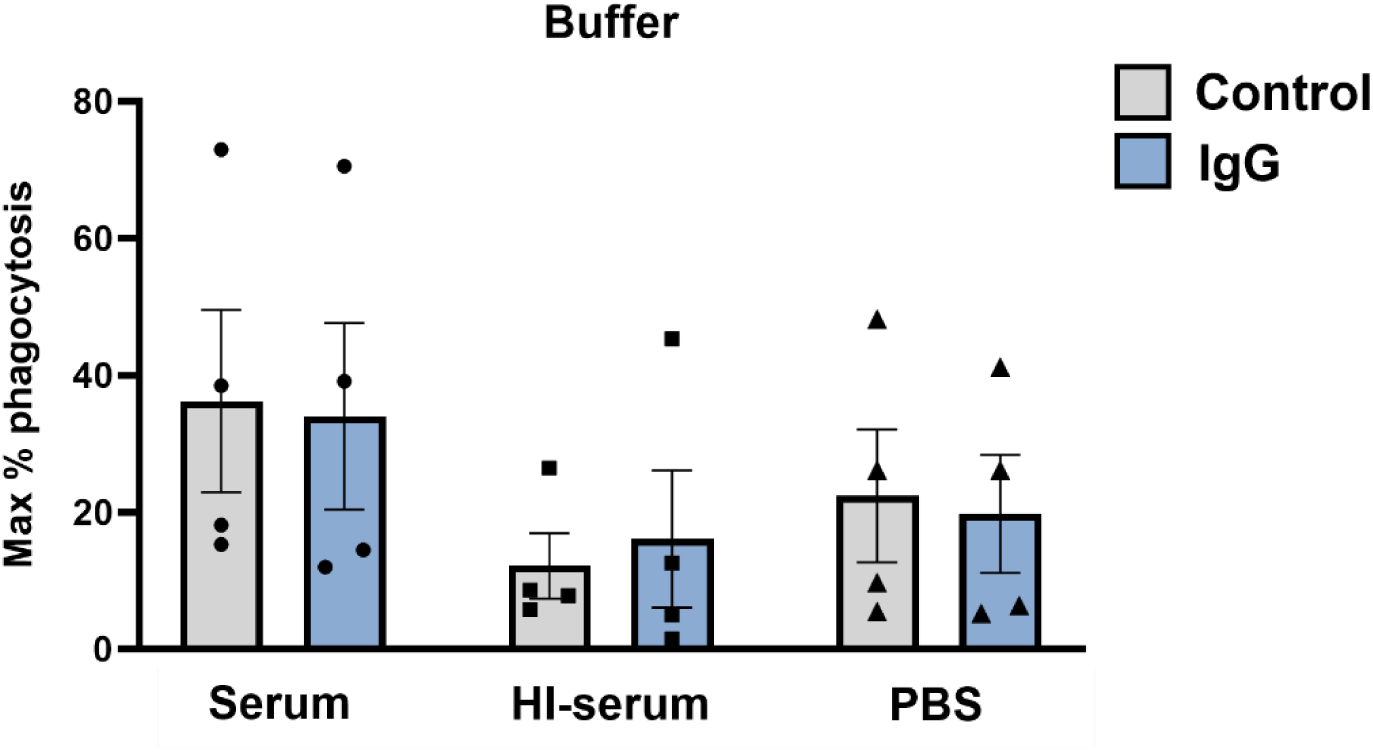
Blocking Fc receptors does not impact myelin phagocytosis. Myelin phagocytosis assays with healthy, human neutrophils (n=4) were performed as previously described. Thereby, human neutrophils were pre-treated for 15 min with buffer (control) or IgG antibodies (100 µg/mL) to block Fc receptors. The percentage phagocytosis was calculated as previously described. Normally distributed data are represented as the mean ± SEM. Data were analyzed using the paired Student’s t-test. IgG: Immunoglobulin G, HI serum: heat-inactivated human serum.

**Supplementary figure 4.**
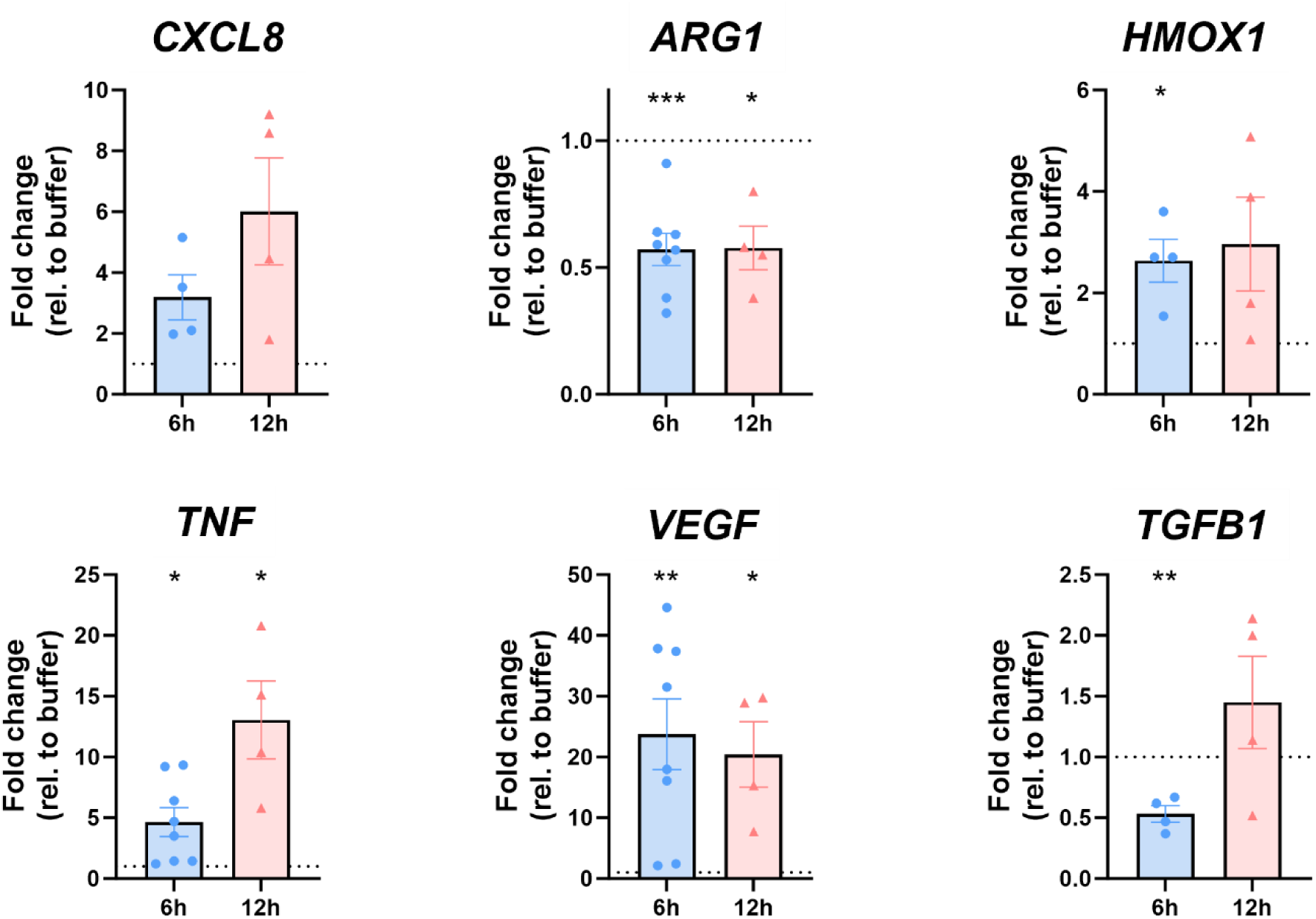
Gene expression levels in human neutrophils after 6 or 12 hours of cytokine stimulation. Human neutrophils (n=4) were incubated for 6 or 12 h with a cytokine mix (50 ng/mL TNF-α, 10 ng/mL IFN-γ and 10 ng/mL IL-1β) or RPMI (buffer). Afterwards, mRNA was isolated and qPCR was performed. Data were normalized to the expression of the housekeeping gene 18S and are represented as fold change relative to buffer-stimulated neutrophils, and are represented as the mean ± SEM. Normally distributed data were analyzed using the one-sample Student’s t-test, with comparison value set at 1; *p<0.05, **p<0.01,***p<0.001.

**Supplementary figure 5.**
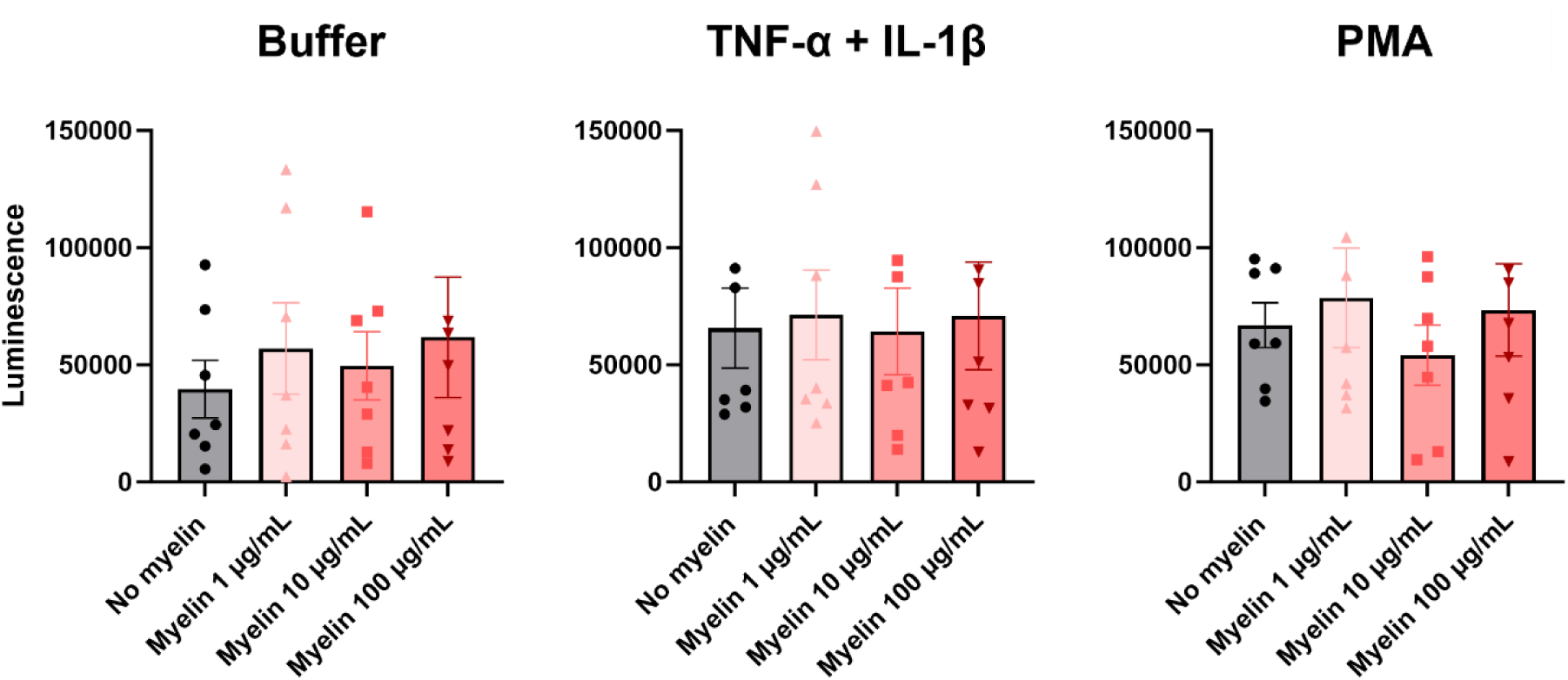
The production of reactive oxygen species by neutrophils after 6 hours of myelin uptake. Purified neutrophils from healthy donors (n=7) were cultured with or without serum-opsonized myelin debris at different concentrations (1, 10 or 100 µg/mL) for 6 hours. Afterwards, cells were harvested, counted and a chemiluminescence-based ROS assay was performed. Therefore, 0.1 x 10^6^ cells were added to each well, and stimulated with RPMI buffer, PMA (150 ng/mL) or a combination of cytokines (priming for 10 min with 10 ng/mL TNF-α and subsequent stimulation with 10 ng/mL IL-1β). Afterwards, luminol (2 mM) was added to the cells, and luminescence was measured for 3 hours at 37°C in a CLARIOstar microplate reader. Maximal luminescence values over time are shown. Normally distributed data are represented as the mean ± SEM. Data were analyzed using a one-way ANOVA.

